# Biomechanics of the extremely elongated neck of the Triassic archosauromorph *Tanystropheus*

**DOI:** 10.64898/2026.06.28.735087

**Authors:** Adam Rytel, Pasha A. van Bijlert, Stephan Lautenschlager, Stephan N. F. Spiekman, Mateusz Tałanda, Tomasz Sulej

## Abstract

Extremely elongate necks have convergently evolved in several amniote lineages, including both aquatic and terrestrial forms (Fig. 1). The development of such a feature brings with it advantages in obtaining food items, but also biomechanical challenges, such as flexibility, stability, lift, and inertia. In *Tanystropheus*, a particularly long-necked Triassic archosauromorph, the neck is composed of only 13, mostly extraordinarily elongated and slender cervical vertebrae and accompanying rod-like, overlapping ribs, making it arguably the most extreme example of neck elongation in tetrapod evolution (Fig. 1;^1–6^). Understanding the function of this remarkable neck provides insights into the limits of neck elongation in amniotes and the evolution of morphological novelties in Triassic reptiles. Here we present the first quantitative biomechanical analysis of the *Tanystropheus* neck using a digital model based on three-dimensionally preserved bones. We assessed its range of motion (ROM) and performed finite element analysis (FEA) on the individual cervical ribs and the neck model in different configurations. Our results indicate that the neck of *Tanystropheus* was not extremely stiff, as previously postulated, and the ribs likely did not impair its movements. They transferred tensile forces towards the base of the neck, similar to what hypothesized for sauropods^7^. This study elucidates the bauplan of an extremely specialized animal and brings us closer to understanding the patterns of achieving neck elongation in vertebrates.

## RESULTS

The reconstructed osteological neutral pose of the neck of *Tanystropheus* (see Methods) is curved, with its central portion located dorsally relative to both the skull and the base of the neck (Fig. 2B). This arch-like reconstruction is further supported by the orientations of the articular surfaces of the centra (Tab. S3).

**Figure 1.**
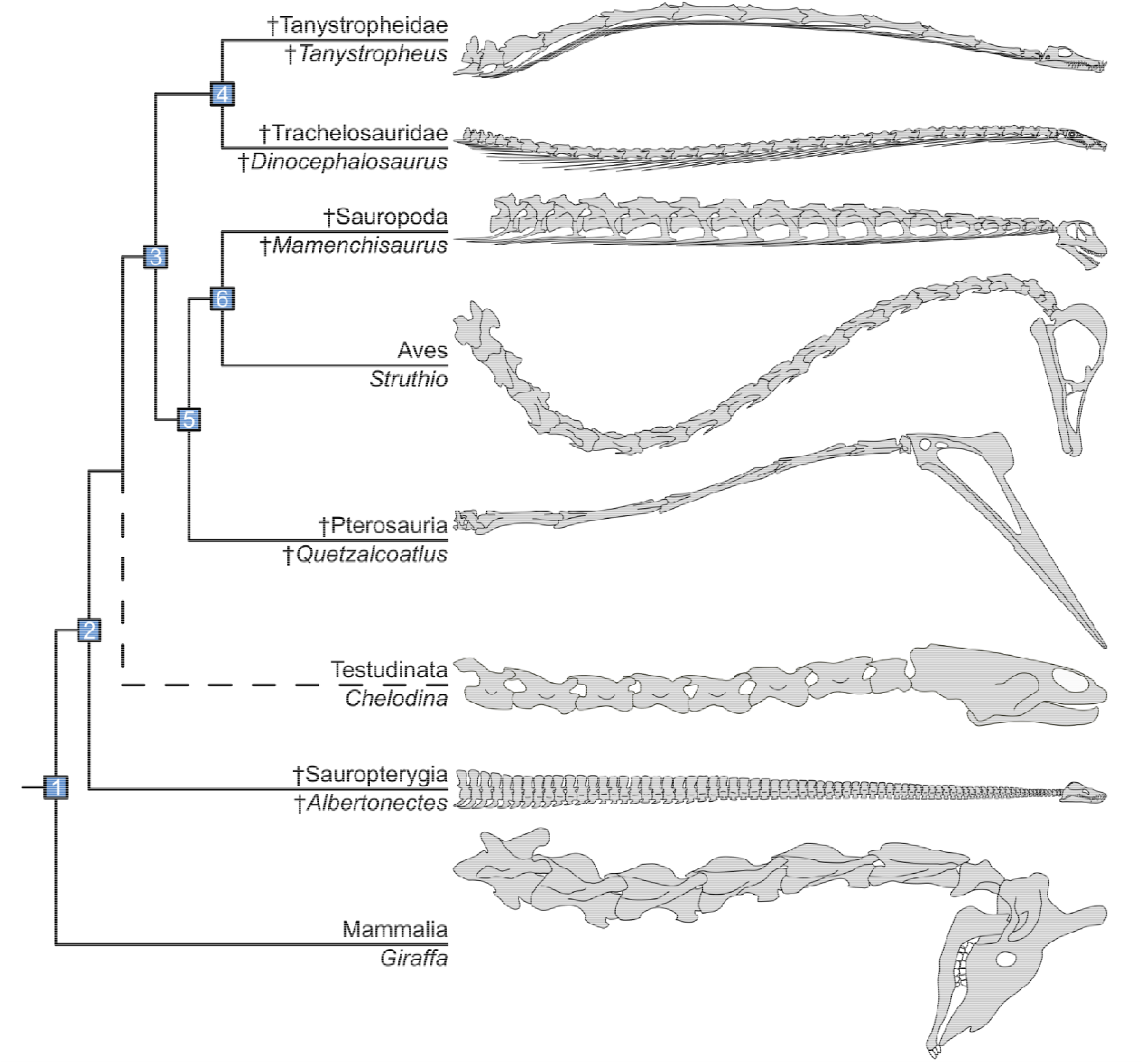
Comparison of the cervical vertebral column anatomy of some long-necked tetrapods. Reconstructions not to scale. Based on: *Tanystropheus conspicuus* – Rytel^6^; *Dinocephalosaurus orientalis* – Spiekman et al.^8^; *Mamenchisaurus youngi* – Pi et al.^9^, Ouyang and Ye^10^; *Struthio camelus* – Mivart^11^; *Quetzalcoatlus* – Padian et al.^12^; *Chelodina longicollis* – Herrel et al.^13^; *Albertonecte vanderveldei* – Kubo et al.^14^, Sachs et al.^15^; *Giraffa camelopardalis* – Badlangana et al.^16^. The numbered nodes correspond to the following taxa: 1. Tetrapoda; 2. Diapsida; 3. Archosauromorpha; 4. Tanysauria; 5. Avemetatarsalia; and 6. Saurischia.

**Figure 2.**
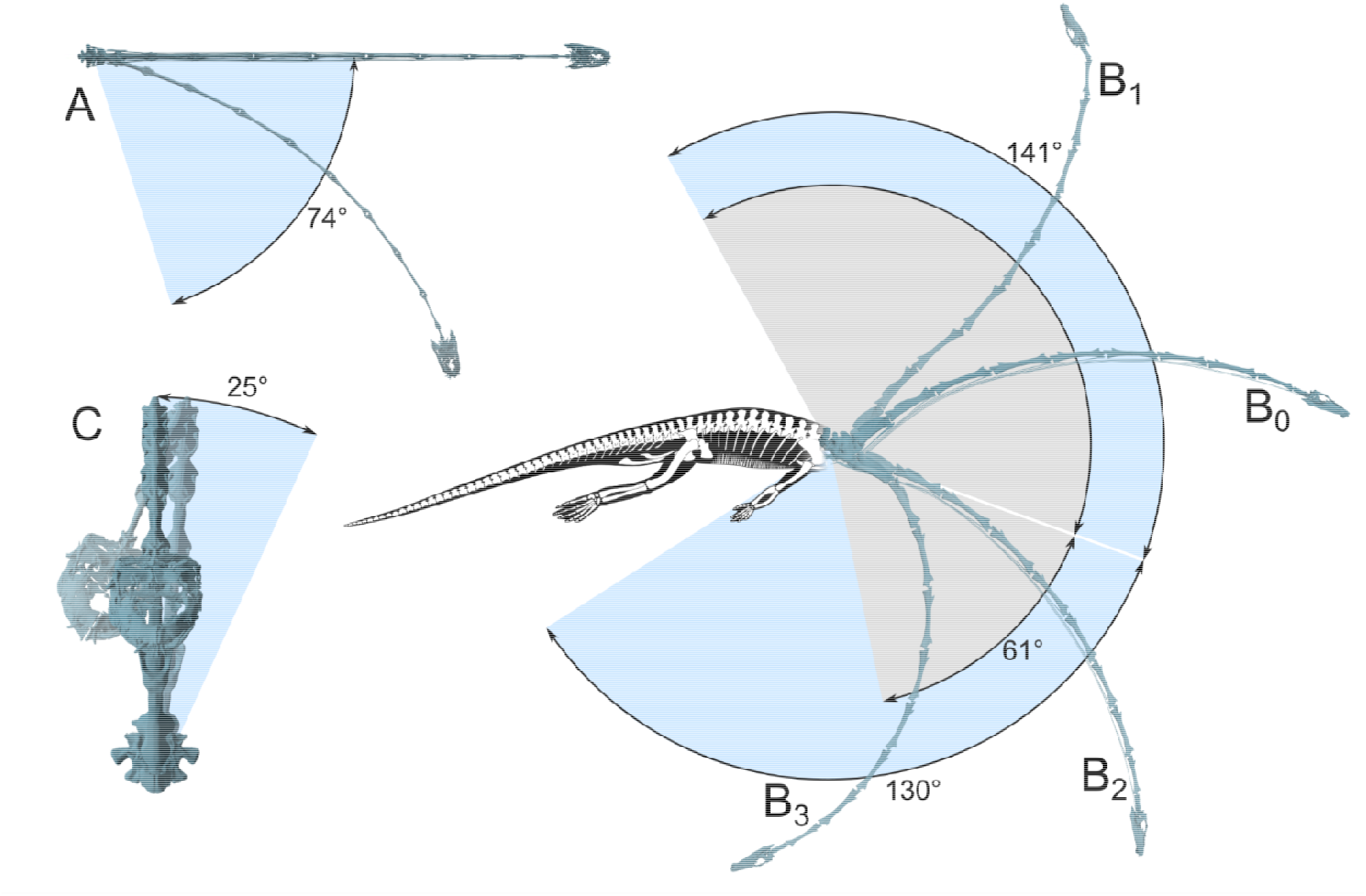
*Tanystropheus* neck postures based on the ROM analysis. (A) Maximum lateral flexion without cervical ribs. (B) Postures within the dorsoventral osteological range of motion: the osteological neutral pose (B0); maximum dorsiflexion (B1) and maximum ventroflexion with (B2) and without cervical ribs (B3). (C) Maximum torsion without cervical ribs. Arcs described between the osteological neutral pose and the maximum flexion (of the skull) in a given plane are indicated in blue for the model including the cervical ribs and in grey for a model without them. All models in orthographic view. *Tanystropheu* reconstruction from Rytel^6^.

The anterior surface of the centrum of the 13th cervical vertebra and the posterior surface of the centra of the 12th and 11th cervical vertebrae were oriented distinctly more dorsally than in the other specimens. In all more anterior cervical vertebrae, the articular surfaces of the centrum are subvertical in lateral view, or slightly ventrally oriented. These characteristics indicate the dorsal inclination of the base of the neck^6,17^, which is also recovered within the model. The middle-anterior portion of the neck is recovered as gradually more ventrally bowing in the anterior direction. In the reconstructed model the distance between the articular surfaces of the articulating middle vertebral centra is considerable, supporting the presence of relatively large amounts of intervertebral tissue^1,18,19^.

The results of the ROM analysis for torsion, dorsiflexion, and lateral flexion are nearly identical for the analytical iterations carried out on the model with and without cervical ribs (Tab. S2). Values for the lateral rotation vary between the joints, but no clear patterns can be noted when comparing between the subregions of the neck. The neck bends laterally in a gentle curve, up to a maximum angle of 73° with ribs and 74° without them (Fig. 2A). Torsion is very limited, with all joints being able to rotate by <5°, which results in a maximum torsion of 25° (Fig. 2C). Maximum dorsiflexion is achieved with the neck being anterodorsally oriented and relatively straight, and the skull pointing posterodorsally. There is a clear disparity between the maximum rotation values reached by the joints associated with the posterior and middle cervical vertebrae, with the former reaching much lower values (2-5°) than the latter (12-17°). The neck is slightly ventrally curved in this posture, with the maximum dorsiflexion reaching 141° for the model with cervical ribs and 142° without them (Fig. 2B). Many articulated specimens of *Tanystropheus* are preserved with a dorsiflexion of the vertebral column (=opisthotonus, see Faux and Padian ^20^; Fig. S2). In the specimens in which the ribs are predominantly still articulated and largely intact, the neck curvature does not exceed the maximum curvature recovered by the ROM analysis.

The maximum ventroflexion achieved by the neck is 130° when the ribs are included and 61° when the ribs are omitted. In both cases, the differences between the rotation achieved by the posterior joints and the anterior joints is opposite to what was noted for dorsiflexion – most of the mobility of the neck during ventroflexion originates from the base of the neck. The middle and posterior cervical vertebrae form distinct modules, mirroring the morphological regionalization reported by Rytel et al.^5^. The rigid ribs nearly completely constrain the movement of the anterior-middle portion of the cervical column. Some posterior vertebrae disarticulate slightly during ventroflexion.

Results of the FEA performed on the cervical ribs individually show that they can bend considerably in all directions before reaching failure stress (Fig. S3). A longer rib can achieve higher curvature at similar stress than a shorter rib. Nevertheless, the curvature of the ribs with peak stresses reaching 100-200 MPa is similar or higher than the curvature of the corresponding neck segments along which they would extend during the maximum rotation in all tested directions of the ROM analysis (compare Fig. S3 and Fig. 2). The bending capabilities of the ribs are further supported by the curvature preserved in a complete left fourth cervical rib of a *Tanystropheus hydroides* specimen PIMUZ T 2819 (see Supp Fig. 2), which is similar to those achieved in the FEA model with peak stresses reaching 100-200 MPa (>70°).

Comparisons of the results from the FEA performed on the neck model with and without ribs reveal a major disparity between them in the compressive/tensile stress distribution pattern (Fig. 3). Without ribs and with dorsally directed force the tensile stress is present across the ventral surfaces of the centrum and the zygapophyses, and the entire dorsal portions of the vertebrae are affected by compressive stress (Fig. 3). The opposite pattern can be observed for the ventrally directed force. With laterally directed force the lateral side of the vertebral column oriented towards it is affected by tensile stress, and the other lateral side by compressive stress (Fig. 3). The inclusion of the cervical ribs into the model affects the distribution of tensile and compressive stresses, but only when the ribs are bound with each other. With ribs treated as individual, non-intersecting elements, stress distribution is nearly identical to the model without ribs. With ribs modeled into bundles, this pattern changes, with a major portion of the stress being transferred from the middle vertebrae (and especially their articular regions) to ribs (Fig. 3). Peak stresses are still observed in the areas around the neural spines of the cervical vertebrae ten and eleven – the only elongate vertebrae of *Tanystropheus* in which the neural spine is well-pronounced along its middle portion and distinctly distally thickened^6^.

**Figure 3.**
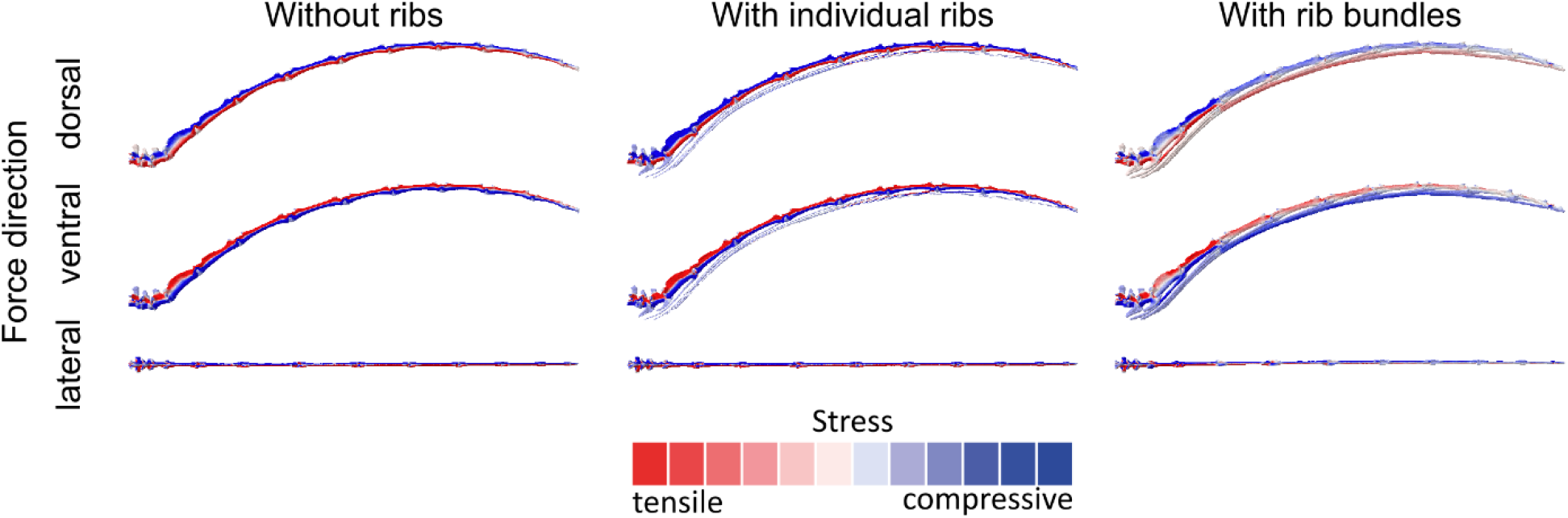
Results of the FEA analysis in models without and with ribs (as either individual elements or bundles), affected by differently directed forces. Colors indicate the tensile/compressive stress distribution, with the areas marked with more intense color being subject to more stress than the areas marked with bleaker colors.

Histological characteristics of the sectioned ribs are fully congruent with the results presented by Jaquier and Scheyer^21^. All analyzed sections show that the shaft was extremely uniform in its internal composition, consisting of avascular compact lamellar bone (Fig. S1). Only the articular heads and the center of the shaft near the articular region were a subject of remodeling.

## DISCUSSION

### Neck of Tanystropheus – stiff or flexible?

Previous studies postulated contradictory hypotheses concerning the mobility of the neck of *Tanystropheus*. Peyer^22^, Wild^1^, and Kummer^23^ hypothesized that it was extraordinarily flexible. Tschanz^18,19^ opposed this view, arguing that the neck of *Tanystropheus* was stiffened by the bundled cervical ribs, which supported the vertebral column and reduced gravitational shearing stresses. In consequence, the neck had to be held horizontally. Renesto^17^ argued that the ribs reduced the neck movements to some extent, but did not render it stiff, as individual thin body rods would be relatively flexible. Nosotti^2^ generally concurred with Tschanz^18,19^, interpreting the stiffening of the neck as an adaptation to cope with underwater propulsion, but noting that an immobile neck would make hunting for nektonic prey difficult. Rieppel et al.^3^ hypothesized that the neck of *Tanystropheus* could be elevated only slightly above the horizontal position, as the ribs severely restricted its movement. The results of the histological study of Jaquier and Scheyer^21^ on the ribs of *Tanystropheus* were also interpreted as indicating a relatively stiff neck.

Results of the ROM analysis provided here indicate that the neck of *Tanystropheus* was not immobile in any direction. The retrieved values are generally similar to those achieved in some directions for other long necked reptiles like plesiosaurs, pterosaurs, or sauropods, despite the comparatively modest number of cervical vertebrae in *Tanystropheus*^12,24–26^. Whereas the aim of this study is to provide general patterns, not definitive values of ROM, and differences in methodology make direct comparisons of the results less feasible, the similarity of the ROM results achieved here for *Tanystropheus* and for the plesiosaur *Cryptoclidus* is particularly intriguing^24^.

With ribs included in the model, only the maximum ventroflexion value decreased substantially. The FEA results indicate that the ribs could bend enough to withstand the stress caused by being bent to the curvature present in the maximal positions of the ROM analysis carried out on the model without the cervical ribs (Fig. 2). Thus, if treated individually, the ribs would likely not restrict the motion of the neck^1,17,18^. As already noted, and valid especially for long-necked taxa, interpretation of results obtained from purely osteological reconstructions not taking into account the effect of soft tissues should be approached with caution^27–30^. Whereas the relatively small number of intervertebral articulations present in the neck of *Tanystropheus* makes this issue less consequential compared to animals with higher cervical vertebral counts, it is difficult to assess how the ribs would behave when bound by interosseous connective tissues and muscles. In articulated *Tanystropheus* specimens the overlapping cervical ribs are preserved in tight association, and likely formed bundles^18^. In life their ability to slide along each other could possibly be limited, together with the total ROM of the neck itself. Nevertheless, a huge part of the maximum ROM, especially during ventroflexion, originates from the base of the neck, which would not be affected by this issue, as the ribs exhibit limited overlap in this section of the column.

The role of the ribs was likely not to brace the neck (see below) and, therefore it still seems unlikely that the hyperelongate morphology of the vertebrae and the associated rod-like ribs stiffened the neck of *Tanystropheus* to the extent proposed by Tschanz^18,19^. The reconstructions presenting the neck as either rather immobile^2,18,19^ or extremely flexible^1,22,23^ are not supported by the results of this study. The neck of *Tanystropheus* could bend in a relatively gentle curve, which would still allow for a considerable reach due its length (Fig. 2). The pose of the neck as restored here is congruent with the insights provided by Wild^1^, Renesto^17^, and Rytel^6^, but in contrast to the hypotheses of Tschanz^18,19^ and Nosotti^2^, who favored a much more horizontal orientation of the *Tanystropheus* neck. The morphology of the posterior cervical vertebrae differs between the species of *Tanystropheus*, with *T. conspicuus* showing more pronounced indicators for the dorsal inclination of the neck than *T. hydroides* and *T. longobardicus*^3,6,17^. The curved posture may have stretched the cervical supraspinous ligaments, storing elastic energy that could have been released during a prey strike lunge^6,18^.

### *Role of the cervical ribs in* Tanystropheus

The hyperelongate, thin and non-bifurcated cervical ribs of *Tanystropheus* are similar to those of the only other tanysaurian that has convergently achieved comparable neck elongation – *Dinocephalosaurus orientalis*^8,31^. In other tanysaurians the ribs can be short and bifurcated (*Trachelosaurus fischeri*), hyperelongate and bifurcated (*Sclerostropheus fossai*), thin and relatively short (*Macrocnemus fuyuanensis*), or thick and short (*Tanytrachelos ahynis*)^32–35^. *Tanystropheus* has proportionally the longest cervical ribs among all known vertebrates, which makes understanding their role in its bauplan particularly interesting. In some specimens the longest individual ribs reach >40% of the combined neck and skull lengths and >20% of total body length (PIMUZ T 2818, see Fig. S2). This exceeds even the ratios estimated for sauropods^36,37^ (e.g., in *Mamenchisaurus* <35% and <20% respectively), for which the role of the cervical rib elongation has been discussed extensively ^7,36,38–41^. The ribs were suggested to provide stabilization for the neck, resistance against torsion, and allowed for a transfer of the cervical musculature towards the base of the neck, which reduces the inertia^36^. Two main hypotheses were formulated – the ventral bracing hypothesis^38,41^ and the tensile member hypothesis^7^. The ventral bracing hypothesis suggests that the ribs supported the neck, transferred compression forces, and counteracted torque. This implies a rather inflexible, horizontally oriented neck, as recovered in the study of Tschanz^18^, who also interpreted the ribs of *Tanystropheus* as transferring compressive forces. In the tensile member hypothesis the tensile forces were transferred instead, and the weight of the neck was reduced by shifting the cervical musculature towards the trunk, allowing for more neck flexibility. In sauropods the ribs are partially composed of ossified tendons, in contrast to *Tanystropheus* in which they are built of avascular lamellar bone^21,39^ (Fig. S1). The longitudinally oriented mineralized collagen fibers, presence of which supports the tensile member hypothesis in sauropods, are absent in *Tanystropheus*^21,39^ (Fig. S1).

Despite the lack of preserved longitudinal mineralized collagen fibers associated with the cervical ribs, our model suggests that the ribs transferred tensile forces, as predicted by the tensile member hypothesis (contra Tschanz^18^). Even though in one of the variations of the model (Fig. 3, ventrally directed force) the ribs are shown as transferring compressive forces, this is highly unlikely, as the hyperelongate ribs would be extremely prone to buckling when loaded in compression. Ventroflexion of the neck would likely require the muscles pulling the ribs, which in turn would incur tensile stress on them. Therefore, it seems more likely that the main role of the hyperelongation of the cervical ribs of *Tanystropheus* was to allow for a shift of the associated musculature towards the torso. This is further supported by the distribution of the muscle attachment sites, which are greatly reduced along the middle portion of the neck, and expanded near its base^1,6,18,19^. Furthermore, two *Tanystropheus* specimens in which the neck was bitten off by a predator or scavenger in a single bite further suggest that the middle portion of the neck was remarkably slender^42^. Finally, the majority of the ribs terminate distally around the same section of the neck, ventrally to the last cervical vertebra. Having long ribs and short tendons and muscles associated with them (instead of short ribs and long tendons) is viewed as an adaptation for a finer control of the neck movements^36^. During movements of a neck with long (or very compliant) tendons, the muscle has to contract a greater distance to also account for the stretch in the tendons. The cervical rib morphology of *Tanystropheus* thus may have functioned more as very stiff tendons, allowing for greater control of the head position, which is necessitated by the extreme length of the neck. The shift of the cervical musculature posteriorly would have also reduced the inertia of the neck. Even considering that the gravitational stress was decreased in an aquatic medium, muscle mass reduction along the neck allowed for its better stabilization, and might have reduced drag during lateral flexion.

### Comparison with other long-necked animals

Although many taxa convergently evolved a long neck, the phylogenetic distribution of hyperelongate cervical ribs among long-necked tetrapods is constrained to archosauromorphs – tanysaurians^8,43,44^ (e.g., *Dinocephalosaurus, Tanystropheus*) and saurischians^36^ (some sauropods and non-avian theropods; see Fig. 1). Their presence seems to be connected with certain specific patterns of neck elongation that appeared both in terrestrial herbivores and aquatic carnivores. *Tanystropheus* is the only known aquatic tetrapod with hyperelongate cervical vertebrae and hyperelongate cervical ribs. *Dinocephalosaurus*, a closely related tanysaurian, had more cervical vertebrae (which were relatively much less elongate than in *Tanystropheus*) than any other archosauromorph and hyperelongate ribs associated with them^8^. Its general bauplan resembled the plesiosaurs. Despite the similarity, in virtually all long-necked sauropterygians the cervical ribs are relatively short, stout, and rarely cross more than one intervertebral articulation. If the presence of elongate cervical ribs is constrained to archosauromorphs, what are the factors controlling their hypertrophy in some long-necked taxa, and atrophy in the others? Even within Sauropoda a major disparity exists in the morphology and length of the cervical ribs between the clades^26,36^, as in the tanysaurians. In long-necked pterosaurs and birds the cervical ribs are extremely reduced (Fig. 1).

Several factors should be considered when interpreting the causes of the observed phylogenetic distribution of the hyperelongated ribs. First, in some taxa the shift of musculature towards the torso might not have been advantageous – in azhdarchid pterosaurs the neck was likely relatively muscular, as it had to support a large head ^45,46^. In all long-necked taxa with hyperelongated ribs the skull is relatively small, presumably due to the effects of gravity and drag (Fig. 1). Whereas in some plesiosaurs the neck to skull length proportions were similar to tanysaurians (Fig. 1), the necks of the former were likely much thicker, which would decrease drag during locomotion and feeding^47^. This is also supported by taphonomic data – despite the rich fossil record of plesiosaurs, there is only one documented case of predatory targeting of their necks^48^, which, in contrast, was documented for two specimens of *Tanystropheus*^42^. This indicates that the neck of the latter was much more slender. Moreover, the short cervical ribs in plesiosaurs served as osteological stops, which might have been energetically more advantageous than using muscle power to keep the neck straight^24^. Finally, the feeding strategy (duration and intensity of the head movements, as well as the neck flexibility required to achieve them) seems to play a major role in the extent of stiffening of the neck tendons ^7,28^. In ostriches, the flexibility of the neck is paramount in obtaining the food items, as the animal tries to reach a large area without moving the trunk, minimizing force and energy requirements during frequent changes of the position of the neck^7^. This is exactly the opposite to what is hypothesized for *Tanystropheus*, which was likely an ambush predator that relied on a limited number of precise strikes^6,49–51^. Thus, the presence and extent of elongation of the cervical ribs in long-necked animals can be correlated with several controlling factors – the relative skull size, the muscularity of the neck, the importance of its flexibility during locomotion and feeding, the preferred posture of the neck, and some other unique characters that are more difficult to assess (e.g., pneumacity). Within the mosaic of bauplans evolved by different long-necked animals, *Tanystropheus* is an important taxon that helps to disentangle the possible connections existing between different aspects of biology of distantly related taxa.

## CONCLUSIONS

This study elucidates the appearance of one of the key anatomical features characteristic to some long-necked animals – the hyperelongated cervical ribs. Whereas they have previously been interpreted as a part of a bracing system that stiffened the neck, our results indicate that at least in *Tanystropheus* the main role of the cervical ribs was instead to transfer tensile forces and shift the mass of the neck towards the torso. This is the first quantitative evidence supporting the tensile member hypothesis, which was formulated for sauropods, but remains difficult to test due to the characteristics of their fossil record^52^. The biomechanical analysis provided here suggests that the individual cervical ribs could bend considerably before breaking. They likely did not restrict the neck movements in *Tanystropheus* in a major way, and could enable finer control over the head movements. This is congruent with the interpretation of *Tanystropheus* as an aquatic ambush predator. These new insights show that despite being composed of a limited number of vertebrae, the neck of *Tanystropheus* could achieve a range of motion comparable to those of some of the long-necked plesiosaurs. This finding rejects previous hypotheses of either extreme stiffness and extraordinary flexibility of the *Tanystropheus* neck, and reveals that this morphological extreme in neck configuration formed a highly effective evolutionary adaptation.

## ACKNOWLEDGMENTS

We wholeheartedly thank K. Przestrzelska, I. Janicki, and P. Bajdek for masterful specimen preparation. We are grateful to Ch. Klug, B. Scheffold, and T. M. Scheyer (all three PIMUZ), E. Mujal and R. Schoch (both SMNS), and J. Kobylińska (ZPAL) for their hospitality and help during the visits and examination of their respective collections. We thank T. Scheyer for allowing us to illustrate some unpublished thin sections of *Tanystropheus* cervical ribs. We thank T. van de Kamp and M. Zuber for their help in CT data acquisition. We want to express our gratitude to the authorities of the Zbrosławice municipality, as well as the community of the Miedary village, especially W. Nawrocki, H. Kupka, I. Hryckiewicz, S. Szczygiel, K. Rode for their long-standing hospitality. We thank K. Pielka, owner of the Miedary claypit area, for support during the excavations. We warmly thank Ł. Czepiński, W. Pawlak, and T. Szczygielski for fieldwork coordination and discussions, M. Dziewiński and J. Jabłoński, for specimen transportation and photography, and all the students and volunteers taking part in the fieldwork in the Miedary site since 2014, particularly those who directly contributed to the discovery of the specimens illustrated in this study: O. Godlewska, A. Kawińska, J. Piwowarczyk, K. Sosnowska, and K. Szuper.

This study was supported by the Polish National Science Centre grants 2017/27/B/NZ8/01543, 2020/39/O/NZ8/02301 (both awarded to T.S.), 2019/35/N/NZ8/03806 (awarded to Ł. Czepiński), and the National Agency for Academic Exchange grant BPN/PRE/2022/1/00009 (awarded to A.R.).

## AUTHOR CONTRIBUTIONS

Conceptualization, A.R., P.A.v.B., and S.L.; methodology, A.R., P.A.v.B., and S.L.; investigation, A.R., P.A.v.B., and S.L.; writing, review & editing, A.R., P.A.v.B., S.L., S.N.F.S., M.T. and T.S.; funding acquisition, A.R., M.T., and T.S.; specimen acquisition, A.R., M.T., and T.S.; supervision, M.T. and T.S.

## INSTITUTIONAL ABBREVIATIONS

PIMUZ, Paläontologisches Institut der Universität Zürich, Zurich, Switzerland; SMNS, Staatliches Museum für Naturkunde Stuttgart, Stuttgart, Germany; ZPAL, Institute of Paleobiology, Polish Academy of Sciences, Warsaw, Poland.

## MATERIALS AND METHODS

### Specimens

To analyze the neck biomechanics of *Tanystropheus* we gathered digital representations (watertight surface models) of the anterior portion of its axial skeleton. These include the skull, cervical vertebrae, the corresponding ribs, as well as the first dorsal vertebra, which originated from two species of *Tanystropheus*. The skull and the atlas-axis complex originated from PIMUZ T 2790, the holotype of *Tanystropheus hydroides* described by Spiekman et al.^50,51^.

To differentiate the anterior, middle, and posterior subregions of the neck we followed Rytel et al.^5^ and Rytel^6^. The models of ribs and most postaxial vertebrae derive from various specimens of *Tanystropheus conspicuus* from the Ladinian site of Miedary, Poland^53^, and were first presented in Rytel et al.^4,5^ and Rytel^6^. They are freely available at MorphoSource: https://www.morphosource.org/projects/000774422. The morphology of the 11^th^ cervical vertebra of *T. conspicuus* was reconstructed based on SMNS 84821 from Schwingen, Germany (see Rytel et al.^4^; https://www.morphosource.org/concern/media/000668709). All mentioned specimens, excluding PIMUZ T 2790, were isolated specimens with excellent three-dimensional preservation. Measurements of the angles between the articular surfaces of the centrum and its ventral margin in lateral view were taken from all postatlantal vertebrae used in the analyses. These data were compared with the curvature reconstructed in the model, to assess if the retrieved posture is supported by the osteological correlates.

Additionally, two cervical ribs of *Tanystropheus conspicuus* from Miedary (the shaft of ZPAL V. 36/1013 and the articular region of ZPAL V. 36/1092) were thin sectioned to gain insights into their internal anatomy. Their histology was then compared with other cervical rib thin sections available for *Tanystropheus* (Supp. Fig. 1).

### Skeletal reconstruction

The anterior portion of the axial skeleton of *Tanystropheus* was reconstructed in Blender 4.5 (https://www.blender.org/). Vertebral models were created by mirroring the better preserved lateral halves of the postatlantal vertebrae along the sagittal plane. Isolated middle cervical vertebrae of *Tanystropheus* cannot be unequivocally referred to a specific position in the vertebral column due to their similar morphology, and no specimen preserves all of them in three dimensions^5,6^. Instead, ZPAL V. 36/1078 was used to represent the cervical vertebrae three to nine (the same model was used in different scales). Shafts of the rib pairs associated with the anterior eleven vertebrae were reconstructed by adding an elongated cylindrical extension to the articular head. The capitulum of the ZPAL V. 36/1000 rib was reconstructed, and the orientation of the capitulum of ZPAL V. 36/1251 was changed to counteract the slight compression of the specimen.

The size of all individual elements was scaled to fit the proportions present in the complete, large individual of *T. hydroides* PIMUZ T 2818 (Supp. Fig. 2), which preserves a nearly complete anterior part of the axial skeleton (see Supp. Tab. 1 for measurements). Measuring the total cervical rib lengths in *Tanystropheus* is difficult due to their fragility and preservation in bundles. In the reconstructed model the rib lengths were adjusted to mirror how many rib pairs are present along different sections of the neck of PIMUZ T 2818, even if the visible elements cannot be directly traced to their articulation with the associated vertebrae.

We analyzed the neck motions with respect to the first dorsal vertebra, which was kept immobile. Positioning of the vertebrae was based on the orientation of the articular surfaces. Only two vertebrae were shown at a time to minimize preconceived notions^26,54^. Each cervical vertebra was positioned so that the articular surfaces of its postzygapophyses are aligned as parallel as possible to the corresponding surface of the prezygapophyses of the more posterior vertebra, and so that the overlap between them is maximized. The position of the vertebra was then further vertically adjusted, to make the corresponding articular surfaces of the centra face each other (to the maximal extent possible without the models intersecting). Ribs were articulated to the vertebrae and the morphology of their shaft was modified to mirror the curvature of the neck. The ribs from one lateral side of the vertebral column were then mirrored to the other. Each succeeding rib pair was positioned dorsolaterally to the pair preceding it. After assembling, the entire model was scaled to match the size of large *T. conspicuus* individuals^6^, from which the isolated elements originated, and measured 325 cm in total anteroposterior length. The resulting model served as the osteological neutral pose for the analyses.

### Rigid kinematic analysis

The ROM analysis was performed with a Blender plugin MuSkeMo^55^ (https://github.com/PashavanBijlert/MuSkeMo), using a rigid body kinematic approach. The aim of this analysis was to recover the general patterns of neck mobility in *Tanystropheus* and assess how it could be affected by the cervical ribs. A center of rotation (joint) was created between each pair of associated centra. Joint placement can affect the outcome of biomechanical analyses^56^ and the chosen locations should be clearly defined. To achieve this, landmarks were added to mark the most anterodorsal, anteroventral, posterodorsal, and posteroventral points on the centra along the neck. The position of the center of rotation was set to the average location of the four mentioned landmarks placed on the corresponding articular surfaces. Joint axes were aligned with respect to each posterior vertebra from the pair as follows: positive x was aligned with the long axis of the vertebra and pointed anteriorly, positive z was oriented mediolaterally and pointed to the right, and positive y pointed dorsally. To achieve this, we fitted cylinders to the vertebral bodies using MuSkeMo’s “Fit cylinder” function, and transferred these orientations to the joints. Bodies were created for each vertebra, and then tied with the joints as the child (preceding) and parent (succeeding) elements, to establish the proximodistal hierarchy of the vertebrae. The atlas-axis complex and the skull were treated as a single rigid element. Two variations of the resulting model were used - with and without the cervical ribs. The ribs were fused with the vertebra, with no movement allowed in the articulation between them.

ROM was checked by employing a script included in MuSkeMo that rotates the entire section of the neck anterior to a given joint around its center of rotation by one degree at a time, until any meshes intersect. Script traversed anteriorly from the most posterior joint (first dorsal vertebra/13th cervical vertebra), recording the last rotation value reached before the intersection for each joint. Maximum ROM was tracked by summing the rotation values reached for the X (torsion), Y (lateral flexion), or Z (dorsoventral flexion) axes of each joint. The maxima were, thereby, the differences in rotation between the osteological neutral pose and the ending pose of the atlas-axis complex and the skull. If the results were asymmetric, the higher maximum ROM value was used.

The reconstructed models of the anterior portion of the axial skeleton of *Tanystropheus* are available as the Supplementary File 1– Blender file with the associated script included.

### Finite element analysis

In order to test the flexibility of the neck, three different biomechanical simulations were performed using FEA: (1) a simulation of an isolated cervical rib to test its bending properties, (2) a simulation of the entire neck with cervical ribs as individual elements or bundles, and (3) the same as two but without cervical ribs. For the FEA of the isolated rib, two straight cervical rib models with different shaft length were tested. For analysis setup and pre-processing, the models were imported into Hypermesh (version 11, Altair Engineering). Solid mesh models were generated from surface models each consisting of ca. 1.5 million elements. In the absence of material properties for reptilian rib bone, properties derived from the biomechanical modelling of human ribs were used^57^. The latter is a common biomedical application of FEA and given the similarity to our study, we used corresponding material properties. To account for the likelihood that the ribs of *Tanystropheus* had somewhat different composition of cortical and trabecular bone, we performed sensitivity tests using a range of material properties (E = 10-20 GPa, □ = 0.1-0.35). As the variation of the material properties did not result in any notable differences in bending behavior, we used the material properties with the highest stiffness (E = 20 GPa, □ = 0.35). These values also fall within the average for most vertebrates^58^. Our results may underestimate the bending capability of the ribs and the flexibility of the neck may have been higher. The rib models were constrained from movement at eight nodes along the articular surface with the corresponding cervical vertebrae. A displacement force was applied to the distal end of the rib models in different directions (dorsal, ventral, and lateral). Results of these analyses were evaluated using overall bending when encountering different von Mises stress thresholds (50, 100, 200 MPa). Given the simplifications inherent in the FEA models, these thresholds represent general indicators of bone stress with bone failure around 200 MPa^59^.

For the two simulations of the full neck model, a surface model of the articulated cervical vertebrae and cervical ribs was imported into Hypermesh. Connective tissues were added in Blender between the centra of successive vertebrae corresponding to the extent of the articular surfaces of the centra. A Boolean operation was applied to remove overlapping surfaces of the connective tissue models with the vertebrae. A single solid mesh was generated for the entire neck model with different material properties applied to the individual components. For the vertebrae and ribs the same material properties as for the simulation of the isolated ribs was selected. For the connective tissue the Young’s modulus was set to 1% of the bone (E = 0.2 GPa, □ = 0.35) similar to the upper range of values for material properties used for sutures studies in lizard biomechanics^60^. Again, this may overestimate stiffness, but as the full neck simulations were performed in a comparative context, material properties were of negligible importance. The model was constrained from movement at 10 nodes at the first dorsal vertebra. The cervical vertebrae were free to rotate and translate about all three axes (six degrees of freedom per vertebra), and these motions were constrained through the material properties of the intervertebral tissue. A load force of 100 N was applied to the second cervical vertebra in different directions (dorsal, ventral, and lateral). This force was chosen to allow for sufficient displacement of the neck, as for this analysis we were interested in the effect of the cervical ribs in a comparative context rather than absolute values. The analyses were performed for each bending direction with and without the cervical ribs attached. Results were evaluated using tensile/compressive stress distribution plots. In contrast to the FEAs for the individual rib models, we did not test for failure stress magnitudes but compared the stress distributions in the models with and without cervical ribs. For the simulation of the rib bundles, individual ribs were aligned so that they are in contact for most of their length. Subsequently, the ribs were remeshed in Blender into a single object that preserved the outline of the ribs but functioned as a single entity for the biomechanical simulations.

