## Supplemental Files (Figures S1-S3 and Tables S1-S3) for "Biomechanics of the extremely elongated neck of the Triassic archosauromorph *Tanystropheus*"

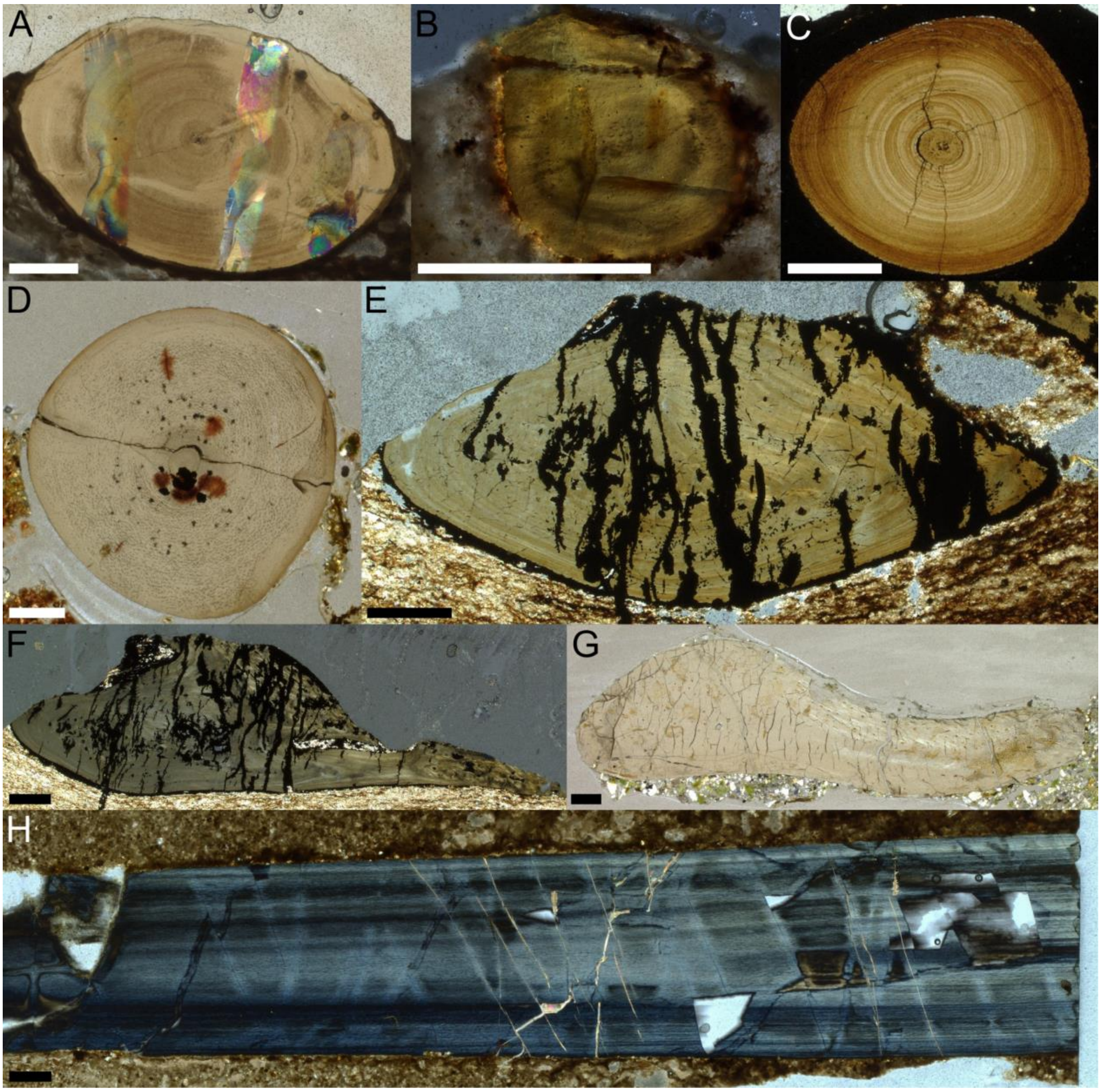


**Figure S1. Histology of the cervical ribs of *Tanystropheus*. Related to Figure 3.** Transverse (A–G) and longitudinal (H) cross sections through the shaft (A–D, H) and the articular portion of the rib (E–G) in normal transmitted (A, C–G) and cross-polarized light (B, H). A, H – SMNS 54057 (*T. conspicuus*). B - PIMUZ T 1277 (*T. longobardicus*). C – PIMUZ T 2789 (*Tanystropheus* sp.). D – ZPAL V. 36/1013 (*T. conspicuus*). E–F – PIMUZ T 2784 (*Tanystropheus* sp.). G – ZPAL V. 36/1092 (*T. conspicuus*). See Wild[1], Jaquier & Scheyer[21], and Rytel[6] for stratigraphic and topographic data concerning the specimens. Scale bars equal 5 mm.

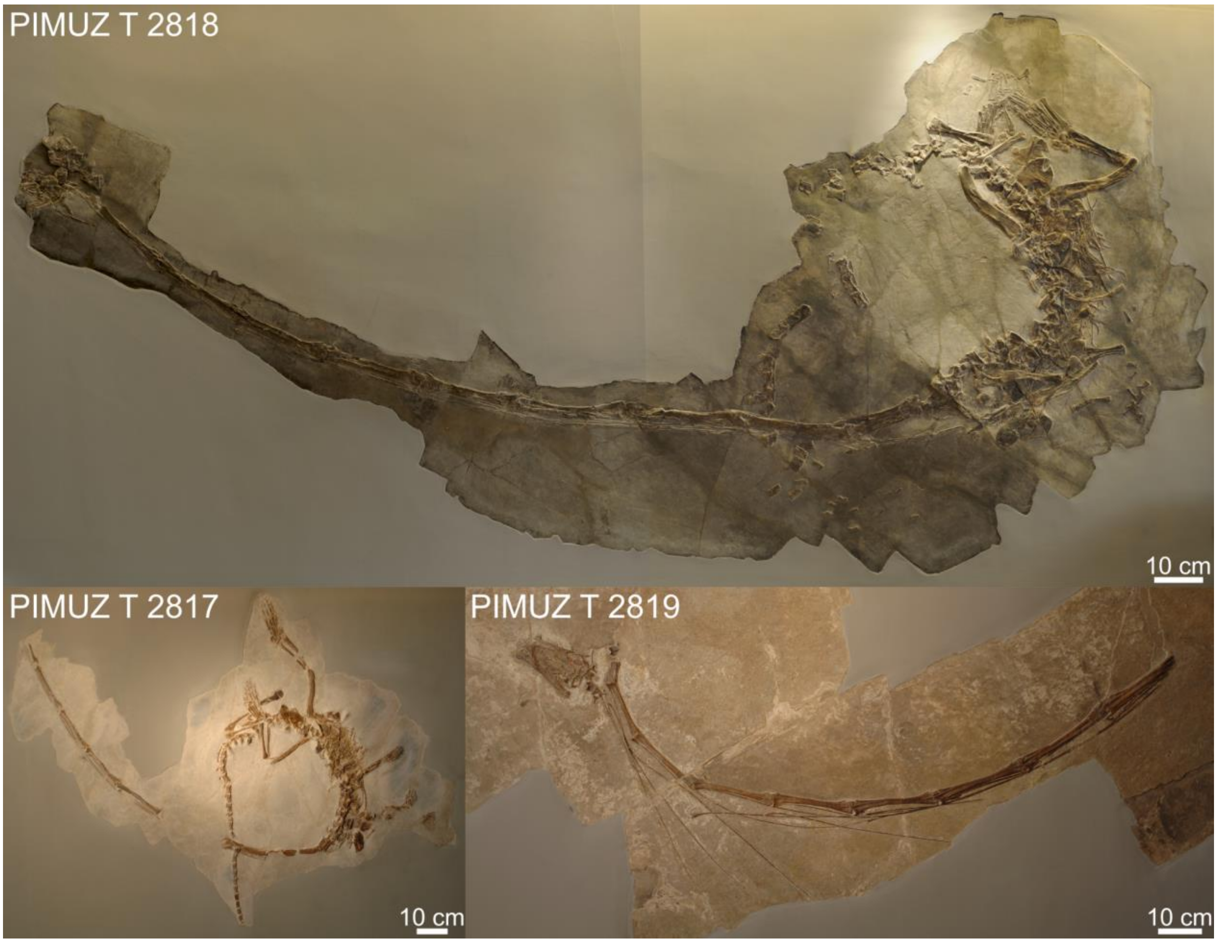


**Figure S2. Articulated specimens of *Tanystropheus* with varying curvature of the neck. Related to Figure 2.**

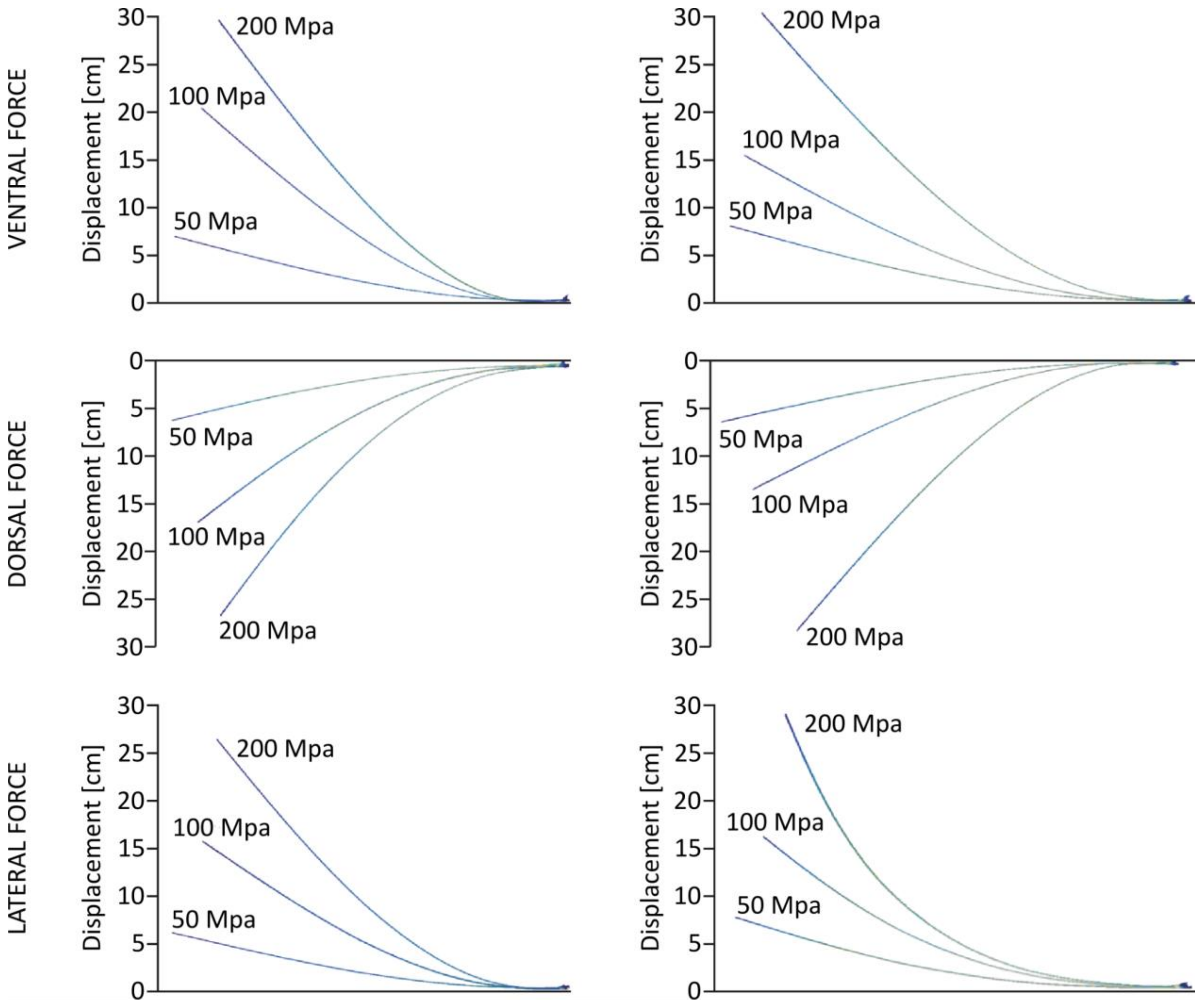


**Figure S3. Results of the FEA analysis performed on the cervical ribs of *Tanystropheus*. Related to Figure 2.** Colors indicate the Von Mises stress distribution [MPa], with the areas marked with brighter color (red) being a subject to more stress than the areas marked with darker colors (blue). White indicates that the region is experiencing stress beyond the current scale.

**Table S1. Measurements of vertebrae and ribs of a large *T. hydroides* individual PIMUZ T 2818 and the corresponding specimens used to reconstruct them within the biomechanical model.** Cervical ribs lengths include only the visible portion of the element (minimal length), usually without the distal end of the shaft.

| Element | Length [mm] | Specimen model used | Notes |
|---|---|---|---|
| CV2 | 68 | PIMUZ T 2790 | |
| CV3 | 170 | ZPAL V. 36/1078 | |
| CV4 | 221 | ZPAL V. 36/1078 | |
| CV5 | 217 | ZPAL V. 36/1078 | |
| CV6 | 230 | ZPAL V. 36/1078 | |
| CV7 | 228 | ZPAL V. 36/1078 | |
| CV8 | 257 | ZPAL V. 36/1078 | |
| CV9 | 246 | ZPAL V. 36/1078 | |
| CV10 | 239 | ZPAL V. 36/186a | |
| CV11 | 178 | SMNS 84821 | |
| CV12 | 78 | ZPAL V. 36/1209 | |
| CV13 | 51 | ZPAL V. 36/2485 | |

| | | | |
|---|---|---|---|
| DV1 | 43 | ZPAL V. 36/1249 | |
| CR2 | 345 | ZPAL V. 36/2433 | Shaft reconstructed |
| CR3 | 609 | ZPAL V. 36/2433 | |
| CR4 | 745 | ZPAL V. 36/2433 | |
| CR5 | 870 | ZPAL V. 36/2433 | |
| CR6 | 954 | ZPAL V. 36/2433 | |
| CR7 | 996 | ZPAL V. 36/2433 | |
| CR8 | 866 | ZPAL V. 36/2433 | |
| CR9 | 737 | ZPAL V. 36/2433 | |
| CR10 | 532 | ZPAL V. 36/2433 | |
| CR11 | 278 | ZPAL V. 36/1019 | |
| CR12 | 96 | ZPAL V. 36/1000 | Capitulum reconstructed |
| CR13 | 82 | ZPAL V. 36/1251 | Capitulum orientation adjusted |

**Table S2. Values of the maximal rotation of the joints achieved in the ROM analysis.**

| Joint | Total rotation [°] | | | | | | | | | | | |
|---|---|---|---|---|---|---|---|---|---|---|---|---|
| | Lateral (left) | | Lateral (right) | | Dorsal | | Ventral | | Torsion (clockwise) | | Torsion (counterclockwise) | |
| | Without ribs | With ribs | Without ribs | With ribs | Without ribs | With ribs | Without ribs | With ribs | Without ribs | With ribs | Without ribs | With ribs |
| DV1/CV13 | 6 | 6 | 6 | 6 | 5 | 4 | 29 | 23 | 3 | 3 | 2 | 2 |
| CV13/CV12 | 10 | 10 | 10 | 10 | 5 | 5 | 16 | 16 | 1 | 1 | 1 | 1 |
| CV12/CV11 | 6 | 6 | 6 | 6 | 5 | 5 | 19 | 14 | 3 | 3 | 2 | 2 |
| CV11/CV10 | 1 | 1 | 0 | 0 | 2 | 2 | 29 | 3 | 0 | 0 | 0 | 0 |
| CV10/CV9 | 4 | 4 | 3 | 3 | 2 | 2 | 5 | 0 | 1 | 1 | 1 | 1 |
| CV9/CV8 | 6 | 5 | 8 | 8 | 17 | 17 | 6 | 1 | 3 | 3 | 2 | 2 |
| CV8/CV7 | 8 | 8 | 6 | 6 | 15 | 15 | 6 | 0 | 2 | 0 | 3 | 0 |
| CV7/CV6 | 8 | 8 | 4 | 4 | 13 | 13 | 4 | 1 | 1 | 1 | 3 | 3 |
| CV6/CV5 | 7 | 7 | 5 | 5 | 15 | 15 | 5 | 0 | 2 | 2 | 3 | 3 |
| CV5/CV4 | 5 | 5 | 4 | 4 | 12 | 12 | 5 | 0 | 2 | 2 | 2 | 2 |
| CV4/CV3 | 8 | 8 | 3 | 3 | 16 | 16 | 4 | 1 | 2 | 2 | 4 | 4 |
| CV3/CV2 | 5 | 5 | 2 | 2 | 35 | 35 | 2 | 2 | 1 | 1 | 2 | 2 |
| Sum | 74 | 73 | 57 | 57 | 142 | 141 | 130 | 61 | 21 | 19 | 25 | 22 |

**Table S3. Differences of angulation of the articular surfaces of the centrum in relation to its ventral edge.**

| Element | Specimen | Centrum articular surface angulation [°] | |
|---|---|---|---|
| | | posterior | anterior |
| CV2 | PIMUZ T 2790 | 96 | 91 |
| Mid CV | ZPAL V. 36 1078 | 97 | 102 |
| CV10 | ZPAL V. 36 186 | 89 | 96 |
| CV11 | SMNS 84821 | 75 | 97 |
| CV12 | ZPAL V. 36 1209 | 72 | 92 |
| CV13 | ZPAL V. 36 2485 | 94 | 73 |
